# Nectar yeast scent additions fail to impact overall bouquet composition and bumble bee visitation in a montane herb

**DOI:** 10.1101/2025.10.17.683152

**Authors:** Nicholas Dabagia, Valerie Martin, Daniel Souto-Vilarós, Robert N. Schaeffer, Rebecca E. Irwin

**Author notes:** Corresponding Author: N. Dabagia. Current Address: Department of Applied Ecology, North Carolina State University, Raleigh, NC 27695 USA.

## Abstract

**Premise of research:** The factors that mediate how foragers locate food supplies are of vital importance in understanding their energy acquisition and survival. Microbes that inhabit floral nectar can play outsized roles in altering nectar chemistry and nutrition, thereby affecting floral visitors.

**Methodology:** Here, we consider one such nectar microbe, the cosmopolitan, specialist nectar yeast, *Metschnikowia reukaufii* (Metschnikowiaceae), by inoculating a nectar analog of the subalpine wildflower, *Corydalis caseana* ssp. *brandegeei* (Fumariaceae), and then characterizing the yeast’s impacts on floral scent composition and the foraging behavior of its main pollinator, *Bombus appositus* (Apidae). We assessed foraging behavior of *B. appositus* in a flower array near the Rocky Mountain Biological Laboratory (Colorado, USA) to test the hypothesis that foragers preferentially visit yeast-inoculated flowers over sterile controls. Additionally, we assessed whether bees spent more time at, and fed more quickly on, yeast-inoculated flowers. In a separate experiment, we tested whether *Corydalis* inflorescences inoculated with *M. reukaufii* had different scent bouquets than sterile inflorescences.

**Pivotal results:** We found that bumble bee pollinators showed no preference for the focal yeast species, with inoculated nectar having no effect on number of flowers visited, time spent on individual flowers, or time spent accessing nectar. Further, the overall scent bouquet compositions of yeast-inoculated *Corydalis* inflorescences were not statistically significantly different than those of sterile inflorescences, despite increased emissions of several volatiles that are known to be produced by *M. reukaufii*.

**Conclusions:** Our findings suggest that *B. appositus* does not respond to the presence of *M. reukaufii* in the nectar of *Corydalis,* and instead, yeast-associated volatile emissions may serve as a reliable cue of a nectar reward that is unused by these pollinators. These findings suggest a few avenues for future research, particularly how morphologically complex, highly scented flowers interact with VOCs produced by nectar-inhabiting microbes, and how floral visitors interpret these signals.

## Introduction

Energy acquisition is critically important for animal survival, growth, and reproduction. While intrinsic factors, such as digestive capacity, can limit energy gain, ecologists are fundamentally interested in the extrinsic environmental factors that constrain energy acquisition (Slobodkin 1962, Wilson 2012, Hall et al. 1992, Kronfeld-Schor & Dayan 2013). A rich body of literature has focused on energy limitations in pollinators, especially bees (Heinrich 1976), which serve as ideal organisms for the study of energy resource limitation given their reliance on sugars from small nectar volumes hidden within flowers (Heinrich & Raven 1972, Heinrich 1975, McCallum et al. 2013). Bees need to visit hundreds to thousands of flowers daily to meet their individual and offspring energetic requirements, with social bees having the additional responsibility of providing for their nestmates (Heinrich 1973, Silvola 1984). Thus, reliable cues of the presence of nectar in flowers may help bees to more efficiently meet their energetic needs.

Bees are known to rely on visual, olfactory, and gustatory cues to identify profitable flowers for foraging (Smith & Raine 2014, Sprayberry 2018, Schaeffer et al. 2019). Recent literature suggests that microbial incidence in floral nectar and their metabolism can significantly alter traits important for visitor localization and assessment of resource quality (Vannette 2020).

Indeed, wildflower surveys have revealed that bacteria and fungi are frequent colonists of nectar (Boutroux 1884, Brysch-Herzberg 2004, Herrera et al. 2008, Herrera et al. 2009, Fridman et al. 2012, Alvarez-Perez et al. 2012, Rering et al. 2024), and that these microbes produce volatile organic compounds (VOCs; Raguso 2008, Wright & Schiestl 2009, Proffit et al. 2020, Souto-Vilaros et al. *in review*) as well as non-volatile compounds, and consume sugars and amino acids (Herrera et al. 2008, Vannette et al. 2013, Vannette & Fukami 2017, Rering et al. 2018). These microbe-mediated changes in nectar phenotype could affect the olfactory and gustatory cues used by pollinators and affect subsequent bee behavior, including changes in visitation rates and time spent feeding on flowers. While microbial scent production in lab-made nectar solutions is relatively well-documented, the effects of microbes on floral volatile emissions and the overall floral-microbe bouquet are less understood (Crowley-Gall et al. 2021). Microbial inoculation experiments in the field have largely looked at floral scent effects of flower-colonizing plant pathogens, showing that infected plants can exhibit altered floral scent profiles (Crowley-Gall et al. 2021). For example, infection with *Erwinia amylovora*, the causal agent of fire blight, led to increased emissions of two green leaf volatiles, 1-penten-3-ol and 3-hexen-1-ol, from apple flowers (Cellini et al. 2019). Plants can respond hormonally and metabolically to microbe-associated molecular patterns of pathogenic, commensal, and beneficial microbes on plant surfaces such as roots, leaves, and flowers (Teixeira et al. 2019). However, we know little about whether nonpathogenic microorganisms, such as nectar specialist microbiota, stimulate changes to plant specialized metabolism in flowers.

Despite documented changes in floral and nectar scent mediated by plant and microbial metabolism (Herrera et al. 2008, Vannette et al. 2013, Vannette & Fukami 2016, Rering et al. 2018, Jacquemyn et al. 2021), studies to date have shown mixed effects on bee foraging behavior (Jacquemyn et al. 2021, Ekemezie et al. 2025). *Metschnikowia reukaufii* (hereafter *Metschnikowia*) is a widespread nectar-inhabiting yeast extensively studied for its tight association with bumble bees as dispersal agents, as well as its influence on nectar traits and bee foraging behavior (Herrera 2014, Rutkowski et al. 2023; Canto et al. 2015, Pozo et al. 2020). Its presence often increases bumble bee visitation, suggesting it can act as a reliable cue of nectar availability by altering VOC profiles through metabolic activity (Schaeffer & Irwin 2014, Schaeffer et al. 2017, Madden et al. 2018, Rering et al. 2018, Yang et al. 2019). For example, Schaeffer et al. (2014) found that two species of bumble bees preferentially visited subalpine *Delphinium barbeyi* (Ranunculaceae) inflorescences inoculated with yeast and probed significantly more flowers on those inflorescences relative to controls. Further, Schaeffer et al. (2017) found that naïve bumble bees preferentially visited artificial flowers inoculated with yeast and spent more time foraging on yeast-inoculated flowers than sterile flowers. Additionally, Yang et al. (2019) found that *B. friseanus* foragers had higher visitation rates on yeast-inoculated flowers of the vine *Clematis akebioides,* leading to increased fruit set in inoculated plants. This preference for yeast-inoculated nectar occurs despite yeast consumption of glucose and sucrose and reducing amino acid content, possibly because consumed yeast cells can provide sterols, lipids, and vitamins (Pozo et al. 2015, Vannette & Fukami 2018, Pozo et al. 2020, Steffan et al. 2019). In contrast, Ekemezie et al. (2025) demonstrated that *B. impatiens* pollinators had no preference for yeast-or bacterium-inoculated artificial flowers over sterile flowers. Similarly, Pozo et al. (2020) found that *B. terrestris* foraging behavior (feeding time and flower choice) on artificial flowers was unaffected by the presence of different yeast species in nectar. Finally, Rutkowski et al. (2022) found that *B. vosnesenskii* workers preferred sterile nectar VOCs to *Metschnikowia-*inoculated nectar VOCs in a laboratory Y-tube olfactometer test. Such variation in bee visitor response, ranging from nectar yeast promotion of flower visitation to deterrence, is likely contingent upon the visitor, microbe, and host plant species identities, in addition to the duration in which microbes are active metabolically in nectar after inoculation (Rering et al. 2018; Schaeffer et al. 2019). Shifts in bumble bee visitation in response to changes in nectar phenotype can have important consequences for host plant reproduction (Herrera et al. 2013; Vannette et al. 2013; Schaeffer & Irwin 2014; Yang et al. 2019). Thus, it is critical to explore how more bee-plant interactions respond to microbial colonization of nectar to gain further insight into their context-dependency.

In this study, we explored how the nectar specialist yeast, *Metschnikowia reukaufii,* affected bumble bee foraging behavior on the flowers of *Corydalis caseana* (hereafter *Corydalis*). First, we quantified fungal density occurrence and density in the nectar of *Corydalis*. Second, using both behavioral assays and gas chromatography mass spectrometry (GCMS), we tested how the presence of yeast-inoculated nectar analogs introduced to inflorescences of *Corydalis* affected bumble bee behavior and floral scent bouquet. Given previous research demonstrating that *Metschnikowia* fermentation alters sugar and amino acid concentrations, resulting in distinct volatile signatures (Madden et al. 2018, Rering et al. 2018), we hypothesized that *Corydalis* inflorescences primed with a yeast-inoculated nectar analog would display a distinct volatile profile and would result in enhanced visitation by its primary pollinator, *B. appositus,* compared to sterile flowers.

## Materials and Methods

### Study system

Research was conducted near the Rocky Mountain Biological Laboratory (RMBL) in Gothic, Gunnison County, Colorado, USA (38°50′N, 106°50′W). We carried out experimental trials in a *Corydalis caseana* ssp. *brandegeei* (Fumariaceae) population in lower Washington Gulch (2940 m, 38.940002, −107.023040), approximately 2 km from the RMBL (Maloof 2000). *Corydalis* is a perennial flowering plant native to subalpine regions of south-central Colorado (Maloof 2000), typically growing in moist, well-drained soil. Mature *Corydalis* have approx. 20 stalks, each containing 5-70 flowers (Maloof 2000). Flowers are bilaterally symmetrical with a single nectar spur of 12-16 mm length (Ownbey 1947). Flowers have a hood made of two dorsiventrally placed petals that cover the flower entrance, mechanically forcing legitimate pollinators through the hood to access the nectar reward and contact plant reproductive organs (Ownbey 1947).

Individual flowers bloom for 4-5 days (Bronstein et al. 2024), with nectar standing crop ranging from 0 to 3.12 μL of nectar per flower (Maloof 2000). Nectar accumulates in flowers over time, with unvisited 5-day old flowers accumulating approx. 6 μl of nectar (Maloof 2000). Species in the *Corydalis* genus have fragrant flowers known to attract bees (Dekebo et al. 2022). In the study region, *Corydalis* is both pollinated and nectar-robbed primarily by bumble bees, with the long-tongued legitimate visitor, *B. appositus* (Apidae) being its dominant pollinator (Wright 1988, Maloof 2000, Heiling et al. 2018).

*Metschnikowia reukaufii* (Metschnikowiaceae) is a specialized nectar-inhabiting yeast, found in multiple animal-pollinated species (Herrera 2014), particularly those pollinated by bumble bees (Herrera et al. 2013, Jaquemyn et al. 2021). It is one of the most abundant yeasts in nectar and is cosmopolitan in distribution (Brysch-Herzberg 2004). *Metschnikowia* is thought to overwinter in bumble bee queens before being dispersed to floral hosts in early spring (Pozo et al. 2018), where it can then be dispersed among flowers and within and among bumble bee colonies and species (Brysch-Herzberg 2004, Pozo et al. 2018). Thus, *Metschnikowia* dispersal is mediated by foraging bumble bees (*Bombus* spp.), other pollinators, and, likely to a much lesser extent, by air, wind, and rain (Martin et al. 2021). Additionally, it is locally abundant in flowers in and around our study site (e.g., Schaeffer & Irwin 2014, Schaeffer et al. 2014), making it an ideal candidate for studying microbial mediation of interactions between bumble bees and *Corydalis*.

### *Enumeration of yeasts in* Corydalis *nectar*

We extracted and pooled nectar from up to six and ten *Corydalis* flowers on an inflorescence in 2021 and 2023, respectively, using Drummond Microcaps capillary tubes (Thomas Scientific, USA). In 2021, we extracted nectar from 31 inflorescences and in 2023, 56 inflorescences.

Nectar samples were diluted in sterile, deionized water and plated on yeast media agar supplemented with the antibacterial compound, chloramphenicol (100 mg L^-1^). Plates were incubated at room temperature (∼20°C) and fungal colonies were counted once growth was visible, yielding the number of colony-forming units (CFUs) per μL^-1^ of nectar (*cf.* Dhami et al. 2018).

### Yeast culture and nectar analog

We cultured *Metschnikowia* collected from *Corydalis* at the lower Washington Gulch site using standard protocols (Dhami et al. 2018, Vannette & Fukami 2018). The *Metschnikowia* strain used in behavior assays was isolated from *Corydalis* nectar in 2022 and used in experiments in 2023. The isolate was identified as *M. reukaufii* by sequencing the D1/D2 subregion of the 26S rRNA gene with NL1 and NL4 primers (*V. Martin, unpublished data,* Kurtzman & Robnett 1998). *Metschnikowia* was cultured weekly from glycerol freezer stocks on yeast media agar plates treated with chloramphenicol.

In behavioral assays, we used a nectar analog that mimics the sugar and amino acid composition of natural *Corydalis* nectar (Souto-Vilaros et al., *in review*), using deionized water with a sucrose concentration of 325.6 g L^-1^ and amino acid concentrations listed in Table S1. We inoculated *Corydalis* nectar analog with a concentration of 1,000 *M. reukaufii* cells μl^-1^, which mimics the natural density of yeast cells within floral nectar (see *Results*, also Herrera et al. 2009). We incubated inoculated nectar at 25°C for 48 h prior to trials to allow cell growth and metabolic activity to modify inoculated nectar. *Corydalis* flowers are in bloom for up to five days (Bronstein et al. 2024), and observed visitation rates (Maloof 2000, Bronstein et al. 2024) suggest a two-day incubation period is within the range of time between initial inoculation by a foraging bumble bee and subsequent visits to an individual flower. This allowed us to standardize the effect of nectar modification by yeast metabolic activity and simulate field-realistic conditions.

### Floral arrays

We conducted choice array trials between July 15 and 25, 2023. The day prior to foraging trials, we bagged *Corydalis* inflorescences with mesh bags to prevent dispersal of microbes to flowers by floral visitors. From the bagged inflorescences on the day of trials, we haphazardly selected six *Corydalis* inflorescences, cut them to a standardized height (∼30 cm), and trimmed them to have 10 flowers per stalk. We then placed the inflorescences in green translucent waterpics in a flight cage (Figure S1) in a 2 x 3 array with 45 cm between waterpics. Three inflorescences each were randomly assigned to yeast-inoculated or control nectar (sterile nectar analog without yeast inoculation) treatments. Prior to applying treatments to flowers, we used 25 μL microcapillary tubes (Thermo Fisher Scientific, Waltham, MA, USA) to remove nectar from each flower. After nectar removal, we injected each flower with 5 μL of the assigned nectar using a 50 μL Hamilton syringe (Hamilton Co., Reno, NV, USA), a volume within the range of nectar volumes accumulating over time in *Corydalis* flowers (Maloof 2000) and could be reliably added with syringes. Syringes were cleaned and sterilized daily using 10% bleach/1% Alconox solution (Alconox Co., White Plains, NY, USA) and UV light; syringes were kept separate and used exclusively for either yeast or sterile treatments.

### Bumble bee observations

Between 08:00 and 10:00 on the day of observations, 12 *B. appositus* workers foraging on *Corydalis* were captured in 50 ml Falcon tubes and starved in a portable cooler on ice for 1 - 2 hours. Bumble bees were introduced individually to the array and allowed time to acclimate to the foraging arena. When the bumble bee began to fly, we allowed her 10 minutes to make an inflorescence choice. Once a bee landed on a flower, we recorded inflorescence treatment, time to feed on individual flowers (assessed by proboscis extension into flower nectar spur), time feeding from individual flowers, and number of flowers probed per inflorescence using a handheld Olympus digital voice recorder (model VN - 541PC). Experimental trials were conducted in observer pairs, with one observer each on either side of the array. We considered a foraging bout over when the bee flew away from the array for a minimum of 2 minutes. When a foraging bout ended, the bee was marked with nontoxic paint (to avoid recapturing; Pocsa, Uni Mitsubishi Pencil, Tokyo, Japan) and released to the area of collection, and any individual flower visited was replenished with 5 μL of nectar according to its treatment assignment. We reused inflorescences between bouts within a day to minimize impact on inflorescence density in the local population. Because reusing inflorescences could affect bee foraging behavior via deposition of bee cuticular hydrocarbons on visited flowers (Witjes & Thomas 2009), we included the reuse of inflorescences across trials as a factor in the analyses (see *Statistical analyses*).

### Floral scent collection and characterization

We compared VOC profiles between *Corydalis* flowers with sterile or yeast-inoculated nectar analog. All open flowers on an inflorescence were treated between 09:00 and 10:30 on July 21 and July 25, 2023 by adding 5 μL of sterile or *Metschnikowia*-conditioned nectar analog (*N* = 8-17 inflorescences per treatment per day). We followed the methods in Campbell et al. (2019) for scent capture, with the exception of the solid-phase extraction (SPE) scent traps (Rering et al. 2020). We prepared SPE adsorbent traps made from Pyrex tubes with a constriction at one end (∼50 mm length, 4 mm outer diameter, 2.4 mm inner diameter; Mountain Glass Arts, NC, USA) with 20 mg of Tenax TA® (60–80 mesh) and 10 mg of Carboxen® 1003 (40–60 mesh; Supelco, PA, USA) secured with silanized glass wool (Sigma-Aldrich, MO, USA) and inert-coated steel mesh (Markes International, UK). Briefly, treated inflorescences were enclosed in a 13 cm x 10 cm nylon bag approximately 2 h after treatments were applied. Headspace volatiles were allowed to equilibrate for 30 min. A scent trap was then inserted into the bag, with the other end connected with tubing to a micro air sampler (Supelco, PA, USA) at a flow rate of 40 mL min⁻¹ for 15 min. Two types of controls - ambient samples consisting of empty bags and vegetative samples of *Corydalis* leaves collected in the same way as floral samples - were collected each day volatiles were sampled.

Analysis of VOCs was completed via thermal desorption and gas chromatography-mass spectrometry (GC-MS) on-site at the RMBL. VOCs were introduced to a Shimadzu GC-MS QP2020 (Shimadzu Corporation, Japan) via a Markes Ultra autoloading system connected to a Markes Unity-xr unit (Markes International, UK). We used ultra-high purity helium as the carrier gas and an Rtx-5MS column (30 m × 0.25 mm internal diameter, 0.25 μm film thickness; Restek, PA, USA). Scent traps were placed within empty inert-coated steel sample tubes (Markes International) and volatiles were thermally desorbed at 220°C for 5 min with a helium flow rate of 75 mL/min. To improve GC-MS detection limits and resolution, volatiles were cryofocused using a cold trap containing 200 mg of Tenax TA (60-80 mesh; Markes International). The cold trap was maintained at −30°C during desorption and then heated to 220°C at a rate of 40°C/min, then held at 220°C for 3 min. GCMS analysis was carried out with the following method parameters: injection temperature 100°C, splitless mode, constant flow of 1.19 mL/min, oven initial temperature 40 °C, hold time 2 min, ramp 5°C/min to 80°C, then ramp 10°C/min to 210°C, then ramp 30°C/min to 275°C (final temperature), for a total run time of 28 min. Ions of sizes 33-300 m/z were measured, with each ion size measured once every 0.3 s. The ion source temperature was 200°C and the interface temperature was 250°C. Peaks were identified and their pseudospectra were compared to the NIST14 mass spectral reference library using the eRah package in R (Domingo-Almenara et al. 2016).

### Compound filtering and verification of database matches

Compounds were retained in the final datasets if a Kruskal-Wallis test found they were significantly more abundant or if they had a mean abundance at least 10 times greater than ambient controls (Eisen *et al*. 2022). In addition, boxplots of each compound by sample type (ambient controls, flowers with sterile nectar, and flowers with yeast-inoculated nectar) were visually examined to ensure that all compounds had higher abundances in flower samples than in ambient controls. Background subtraction was performed by subtracting the mean ambient control abundance from the abundances of each compound across samples. Kovats retention indices (RIs) were calculated using a series of n-alkane standards. RIs were used to verify initial database identifications. Database identifications were further verified by comparison to retention times of analytical standards when available. Compound identities not authenticated with a standard were recorded as tentatively identified. Compounds were labeled as unknown if database matches were poor and identities could not be confirmed with analytical standards. The ten most abundant ions for unknowns are listed in the supplemental information (Table S2).

### Statistical analyses

All statistical analyses were conducted in R 4.4.3 (Trophy Case; R core team 2025). Data manipulation and visualization were performed using the *tidyverse* collection (Wickham et al. 2019).

### Floral arrays

A total of 62 bumble bees were tested, of which 23 made a choice and visited at least one flower within 10 minutes of beginning to fly within the array. Thus, our statistical analyses focused on these 23 bees. Due to the strongly skewed nature of the data, we used one-sided, non-parametric Wilcoxon Rank Sum Tests (R: wilcox.test, package *stats* (R core team 2025) to test the hypotheses that bumble bees exhibited higher median foraging times per flower, faster median time to feed on flowers, and higher median numbers of flowers visited in the yeast-inoculated treatment relative to control treatments. Because we reused flowers between bumble bee replicates within a day, we also tested for the effect of previous bee visitation on subsequent bee foraging behavior with Wilcoxon Rank-Sum Tests.

### Floral scent

A PERMANOVA was performed to test for significant effects of *Metschnikowia* inoculation on Bray-Curtis distances of flower scent compositions (R: distance, package *phyloseq*; adonis2, package *vegan*) (McMurdie & Holmes, 2013; Oksanen *et al.,* 2024). Compounds with significant differential abundances between sterile- and *Metschnikowia-*treated inflorescences were detected using the DESeq2 package in R (Love *et al.,* 2014).

## Results

### Yeast abundance in Corydalis nectar

In both 2021 and 2023, many nectar samples from *Corydalis* inflorescences contained culturable yeasts: 61% (19/31) of samples in 2021 and 48% (27/56) of samples in 2023. In flowers that contained yeast, concentrations were as high as 3,845 colony-forming units (CFU) per μL nectar in 2021 and 7,450 CFU per μL in 2023 (Fig. 1).

**Figure 1:**
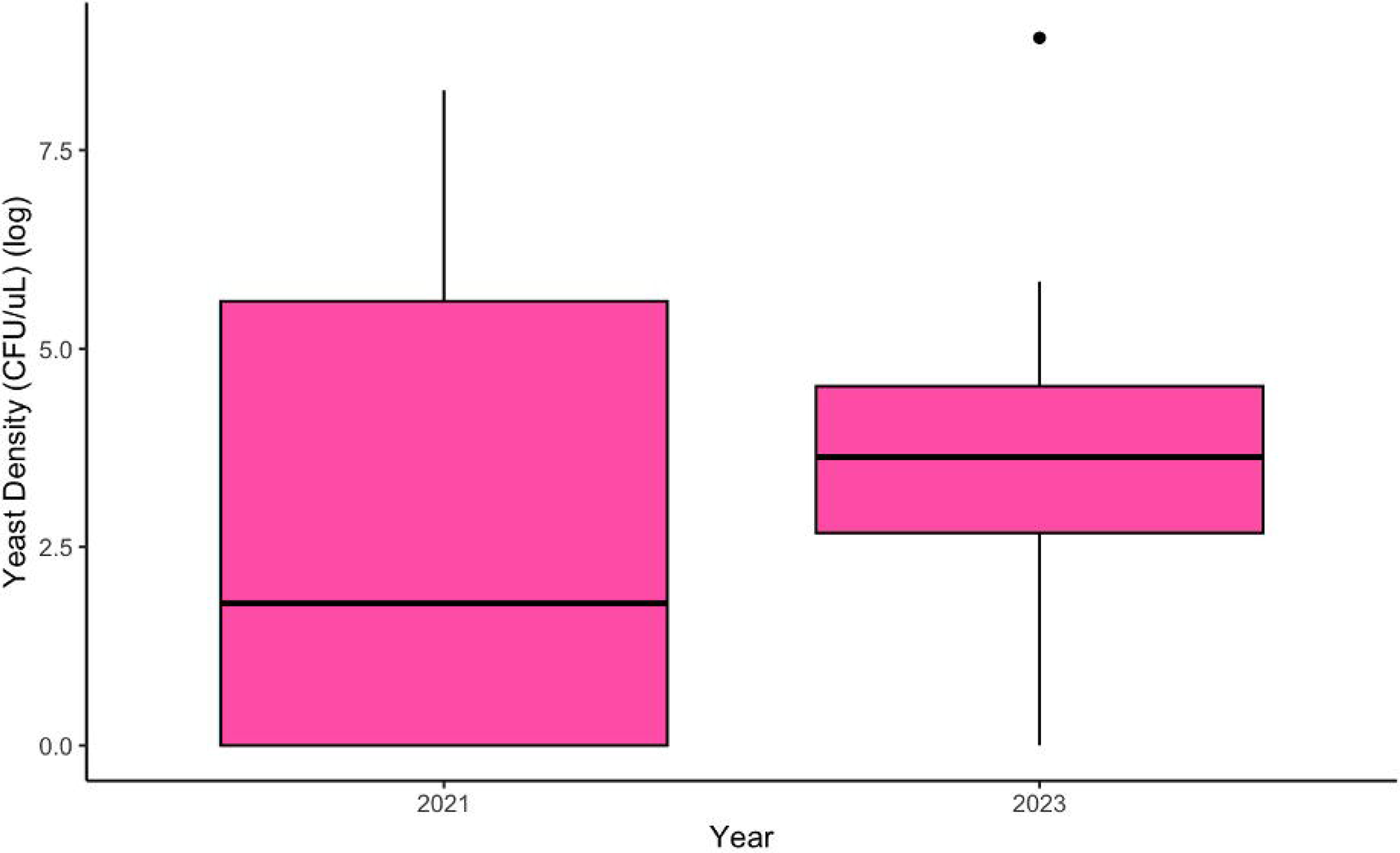
Boxplot of log yeast density in *Corydalis* nectar in 2021 and 2023 at the Washington Gulch population. The boxplot of yeast densities includes samples with and without yeast. Boxplot showing densities of yeast across two years of sampling. 2021 shows a box with mean ∼log(2.0), ranging from log(0) to ∼log(8). 2023 shows a box with mean ∼log(3), ranging from log(0) to log(5.5), with a single outlier at log(8.5).

### Floral arrays

We found that *B. appositus* workers showed no preference for and had no differences in foraging behavior at yeast-inoculated vs. control flowers of *Corydalis*. Of the 23 bees that made a foraging choice, 20 individuals visited at least one sterile flower, 18 visited at least one yeast-inoculated flower, and 12 visited at least one sterile flower and at least one yeast-inoculated flower.

Our trials revealed that *B. appositus* workers visited similar numbers of yeast-inoculated and control flowers per foraging bout (Fig. 2A; W = 185, p = 0.45, one-sided, n _yeast bouts_ = 18, n _sterile bouts_ = 20), spent similar amounts of time feeding at yeast-inoculated and control flowers (Fig. 2B; W = 21202, p = 0.91, one-sided, n _yeast flowers visited_ = 197, n _sterile flowers visited_ = 237), and did not differ in the time it took them to feed on flowers of the two treatments (Fig. 2C; W = 16296, p-value = 0.61, one-sided, n _yeast flowers visited_ = 197, n _sterile flowers visited_ = 237). Across all bouts, bumble bees visited a total of 197 yeast-inoculated flowers and 237 sterile control flowers. Bees spent a median time of 4.2 seconds visiting yeast-inoculated flowers over 197 flower visits with a range of 0.5 to 66.8 seconds, while they spent a median time of 4.9 seconds on control flowers over 237 flower visits, with a range of 0.5 to 76.1 seconds. Finally, bumble bee foragers began to feed in median 0.6 seconds with a range of 0.2 to 7.4 seconds on control flowers, and median 0.5 seconds with a range of 0.2 to 4.1 seconds on yeast-inoculated flowers.

**Figure 2:**
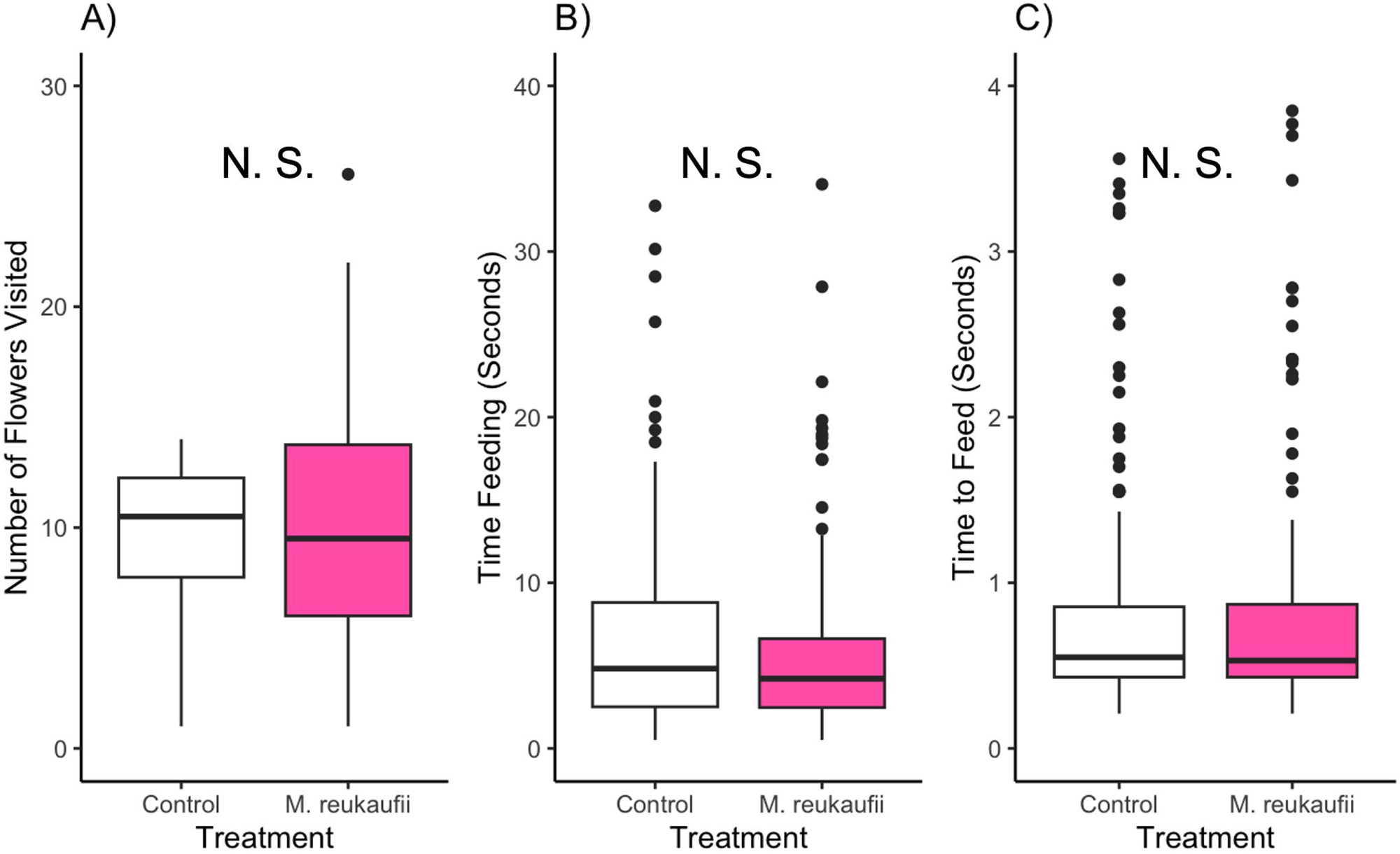
Boxplots of bumble bee foraging to yeast-inoculated (pink) or control (white) flowers: **A)** Number of flowers per foraging bout visited by *B. appositus*, **B)** Time (seconds) *B. appositus* spent feeding per flower, and **C)** Time (seconds) it took *B. appositus* to feed. N.S. indicates no significant difference in metrics of bumble bee foraging between control and *Metschnikowia*-inoculated flowers. Panel with three boxplots showing insignificant relationships between metrics of bumble bee foraging (number of flowers visited, time spent feeding per flower, time to begin feeding per flower) and yeast-inoculated or sterile nectar. There are several outlying points for both sterile and inoculated boxplots for time feeding per flower and time to feed per flower.

Turning to the issue of reusing stalks among trials within a day, we found that previous visitation did not significantly affect either number of flowers visited per bout (W = 172.4, p = 0.9033, two-tailed, n _bouts to visited flowers_ = 14, n _bouts to unvisited flowers_ = 24) or time spent foraging (W = 20198, p = 0.4214, two-tailed, n _visits to visited flowers_ = 153, n _visits to unvisited flowers_ = 277). However, foragers did feed significantly faster on flowers that had been previously visited (0.5 seconds, median time to feed on visited flowers, range = 7.1 seconds, median time to feed on unvisited flowers = 0.6, range = 3.9 seconds; W = 18094, p = 0.01237, one-sided, n _visits to visited flowers_ = 141, n _visits to unvisited flowers_ = 225).

### Floral scent

Across both yeast-inoculated and control flowers, 180 individual volatile compounds were detected. The floral scent composition was dominated by monoterpenes (25 compounds), fatty acid derivatives (20 compounds), and benzenoids (17 compounds). Beta-myrcene (29.6% of average total peak area), phenylethyl alcohol (18.0%), limonene (15.9%), 2-cyclopentylcyclopentanone (5.4%), phenylethyl acetate (5.2%), and beta-thujene (3.2%) were the most abundant compounds in the floral scent profiles.

Detected *Metschnikowia* volatiles included isoamyl alcohol, active amyl alcohol, ethyl acetate, isobutyl acetate, active amyl acetate, hexyl acetate, isovaleraldehyde, phenylethyl alcohol, and phenylethyl acetate. Four of the known *Metschnikowia* volatiles had significantly higher emissions from inflorescences treated with *Metschnikowia-*inoculated nectar: ethyl acetate, isobutyl acetate, isovaleraldehyde, and active amyl alcohol (Figure 3). Moreover, a total of 35 compounds had significantly higher emissions from *Metschnikowia*-treated flowers, while six compounds exhibited lower emissions (Figure 3). Despite these measured differences for individual compounds, a PERMANOVA analysis did not detect a significant difference overall in the floral volatile compositions of *Corydalis* inflorescences inoculated with *Metschnikowia* compared to those of the sterile inflorescences (F_1,24_ = 1.245, P = 0.249).

**Figure 3:**
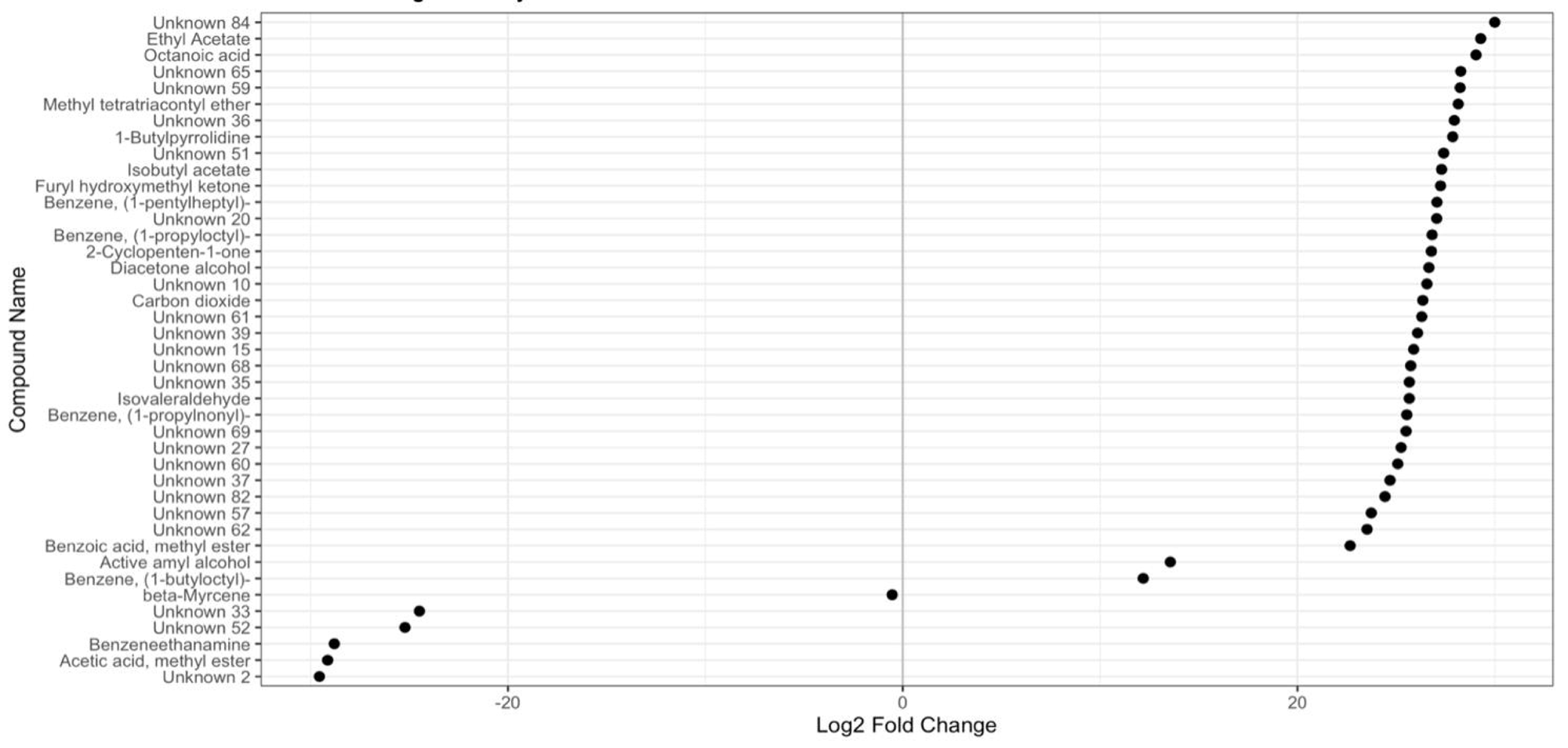
Volatile compounds with significantly different emissions from *Corydalis caseana* ssp. *brandegeei* inflorescences treated with control or *Metschnikowia*-fermented nectar. Volatile compounds with significantly different emissions from *Corydalis caseana* ssp. *brandegeei* inflorescences treated with control or *Metschnikowia*-fermented nectar.

## Discussion

Here, we tested the hypothesis that microbes present in nectar play a role in bee foraging behavior, providing a cue of the presence of nectar in flowers. We found that *Corydalis* flowers commonly contain yeasts, which were present in about half of flowers sampled in two years.

However, we found that our focal yeast species, *M. reukaufii*, had no significant effect on the foraging behavior of *B. appositus* workers at *Corydalis* flowers, indicating that the nectar yeast did not provide a cue of nectar availability used by *B. appositus* pollinators. Furthermore, *Corydalis* flowers did not have a detectable difference in overall flower scent bouquet when treated with yeast-inoculated compared to sterile nectar analog, despite individual compound emissions being affected by *Metschnikowia* presence and metabolism. Taken together, our results suggest that *B. appositus* workers do not prefer or avoid *Corydalis* flowers containing *Metschnikowia*-inoculated nectar, and lack of overall floral scent bouquet between the two treatments may be a driving factor.

Prior studies of the impacts of nectar yeasts on the behavior of floral visitors have not found a consistent pattern of the effect of nectar yeast, suggesting that plant-pollinator-yeast interactions are context-dependent. Some studies have found a positive effect of nectar-yeast inoculation (Herrera et al. 2013, Schaeffer et al. 2014, Schaeffer et al. 2017), including on the foraging of nectar-robbing *B. flavifrons* and *B. bifarius* workers foraging from *Corydalis* flowers (Souto-Vilaros et al., *in review*), while other studies have found a null or negative effect of yeast inoculation on bee foraging behavior, including on artificial flowers (Pozo et al. 2020, Ekemezie et al. 2025). In another group of bee foragers, honey bees were not affected by the presence of yeasts in nectar when individuals encountered them in laboratory assays (Rering et al. 2021), synthetic flowers (Good et al. 2014), or *Asclepias syriaca* flowers (Kevan et al. 1988). Like these latter studies, we found that *B. appositus* workers did not respond to the presence of *Metschnikowia* in *Corydalis* flowers. There was no significant difference in any metric of bee foraging behavior between yeast-inoculated and sterile nectar in *Corydalis* flowers. Taken together, we suggest that future work should assess the impact of nectar yeasts on bumble bee behavior in a wider range of flowering species.

We had predicted that any differences in bee foraging behavior would be driven by VOC profile differences between yeast-inoculated and sterile control flowers. Bees are known to detect and respond to volatiles emitted by flowers, including several of the volatiles identified in the *Corydalis* scent bouquet (Dötterl & Vereecken 2010). Interestingly, while Souto-Vilaros *et al*. (*in review*) found that *Metschnikowia-*inoculated *Corydalis* nectar analog emitted several yeast-associated volatiles, we found no significant difference in floral scent compositions between the flower treatments in *Corydalis*. Our results suggest that, while four yeast-associated volatiles and several plant-produced compounds were emitted from *Metschnikowia-*inoculated flowers in greater abundances (Figure S2), these changes were not enough to significantly shift the overall scent profiles. Past studies of flower yeasts have focused on volatiles emitted from synthetic nectar-like media (Rering et al. 2018, Rering et al. 2021, Sobhy et al. 2025), rather than *in planta*, as we did. *Metschnikowia* species have been shown to significantly alter the scent of several types of nectar analogs (Schaeffer & Irwin 2014, Schaeffer et al. 2017, Madden et al. 2018, Rering et al. 2018, Yang et al. 2019, Souto-Vilaros et al. *in review*). We aimed to determine whether yeast-produced nectar compounds were detectable among the scent compounds produced by the odiferous *Corydalis* flowers and whether inoculation with yeast resulted in changes to the plant-produced compounds. We were able to detect *Metschnikowia* volatiles in the headspace of *Corydalis* flowers, though these volatiles combined contributed less than 1% of the total floral scent emissions. The six most abundant compounds emitted by *Corydalis* inflorescences, constituting 77.3% of the total volatile emissions, were plant-produced. Of these six compounds, phenylethyl alcohol and phenylethyl acetate are known to be produced by both flowers and yeasts; however, the emission rates of these two compounds indicated that they were plant-produced: these two benzenoid compounds are emitted from nectar by *Metschnikowia* in smaller abundances than active amyl alcohol (Rering et al., 2018), which was a minor constituent of the floral scent bouquet and only detected in *Metschnikowia*-inoculated samples, contributing 0.03% of average total peak area. If *Metschnikowia* was the main producer of phenylethyl alcohol and phenylethyl acetate, they would have been detected in much smaller quantities, indicating that *Corydalis* emitted the large observed quantities of these two benzenoid compounds. We did not detect significant changes to the floral scent bouquet, indicating that there was no effect or limited effects of *Metschnikowia* on the specialized metabolism of *Corydalis* flowers.

We cannot rule out the possibility that the complex morphology of *Corydalis* flowers may be responsible for driving the lack of effect of yeast on bee foraging and VOC profiles. *Corydalis* flowers have a complex morphology, with hooded flowers and long nectar spurs (Maloof 2000). The hooded flower morphology and nectar spur could have limited the diffusion of nectar yeast VOCs from the flower. Indeed, we found that *Metschnikowia-*inoculated nectar in tubes placed next to flowers emitted higher amounts of yeast-produced volatiles than the same amount of nectar placed near nectaries within flowers, as in this experiment (V. Martin, *unpublished data*). The role that flower morphology plays in microbial volatile diffusion and transmission is poorly understood (Adler et al. 2021), although recent work has shown that different surfaces on complex flowers can have different signatures of VOCs (Heuel et al. 2025). Similarly, Souto-Vilarós et al. (*in review*) found that bumble bee nectar robbers located artificial robbing holes along the nectar spur faster in yeast-inoculated flowers than sterile controls, suggesting that robbing holes may enhance the diffusion of VOCs from within the flower. In addition, complex flower morphologies require learning and handling time by bees (Laverty & Plowright 1988, Laverty 1994, Gegear & Laverty 1995). Yeast metabolism in nectar has known gustatory effects on bees (Herrera et al. 2013, Schaeffer et al. 2017, Schaeffer et al. 2019). It is possible that by the time *B. appositus* recognizes potential differences in scent and taste between nectar with and without yeast in *Corydalis*, they have already committed to foraging on that flower (Krishna & Kaesar 2018) and opt to do so regardless of the yeast and its activity in the nectar. Future research is needed to understand how flower morphology affects the transmission of microbial signals from floral nectar to floral visitors.

We found that *Metschnikowia* inoculation of *Corydalis* flower nectar did not significantly impact the overall VOC bouquet of inflorescences or the foraging responses of bumble bees when they encountered inoculated flowers in a floral array. Because of the varied responses that pollinators may have to microbe-inhabited nectar, and the impacts of this altered behavior on pollinator and host plant fitness, it is vital that we continue to characterize and understand these context-dependent and piecewise interactions between nectar microbes, their floral hosts, and floral visitors. We suggest that future research takes aim at understanding how flower morphology impacts the production, emission, and perception of both floral and microbial volatiles.

## Acknowledgements

We thank the RMBL and Gunnison National Forest for access to field sites. Jennifer Reithel was invaluable in facilitating the logistical aspects of field and laboratory work at the RMBL, and Dylan Kitselman prepared the borosilicate glass tubes for our scent traps. We thank *Corydalis caseana*, *Bombus appositus*, and *Metschnikowia reukaufii*, the land that sustains them, and the Ute peoples who are the original stewards of the lands now known as Colorado. This work was funded by National Science Foundation grants DEB-2211232 to RS, DEB-2211233 to RI, and DBI-1624073 to RMBL. VM acknowledges fellowship support from Utah State University, the Rocky Mountain Biological Laboratory, the Mycological Society of America, the Colorado Native Plant Society, the Botanical Society of America, the Garden Club of America, and Graduate Women in Science. Any opinions, findings, conclusions, or recommendations are those of the authors and do not reflect the views of the funding agency or funders.

