## Supplementary Information for "Nectar yeast scent additions fail to impact overall bouquet composition and bumble bee visitation in a montane herb"

**Supplemental Information**

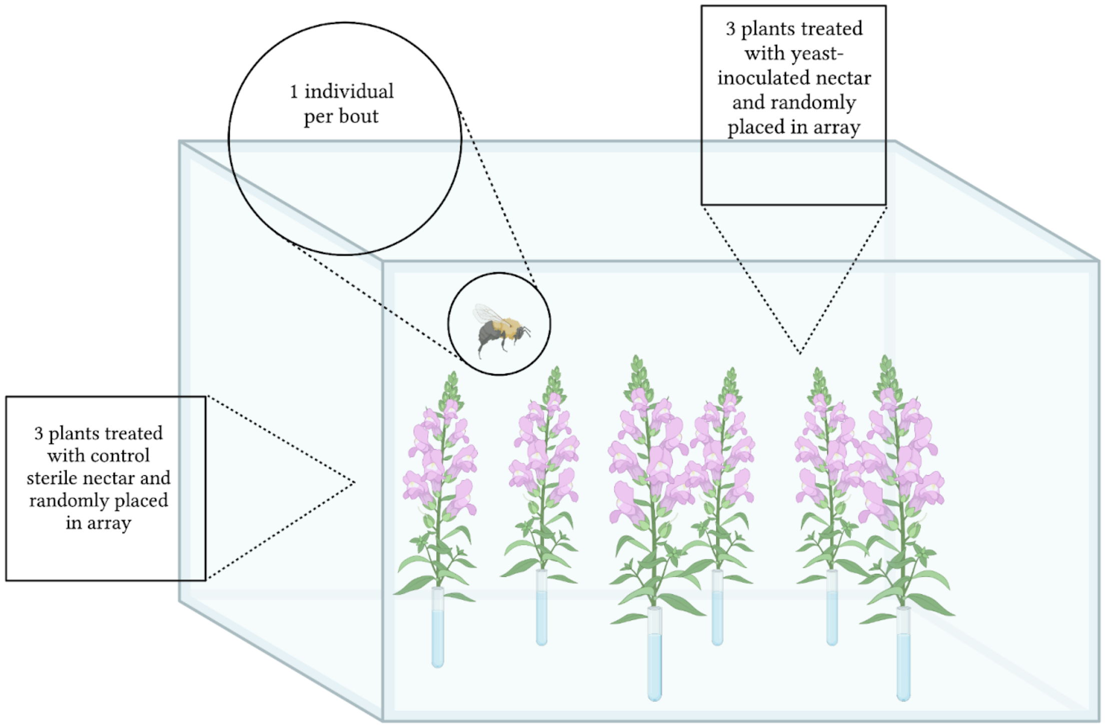


**Figure S1.** Experimental array in a flight cage for bumble bee behavioral trials.

**Table S1.** Table describing concentrations of amino acids (mg/L) in *Corydalis caseana* spp. *brandegeei* nectar. These concentrations of amino acids were used to make *Corydalis* nectar analog for bumble bee behavior trials.

| Amino Acid | Corydalis mg_L |
|---|---|
| **Aspartate** | 2.383525222 |
| **Glutamate** | 8.918193778 |
| **Asparagine** | 4.110364889 |
| **Serine** | 5.446749111 |
| **Glutamine** | 4.895405778 |
| **Histidine** | 4.895405778 |
| **Glycine** | 3.381586111 |
| **Threonine** | 0.9187495 |
| **Arginine** | 0.9187495 |
| **Alanine** | 2.946735 |
| **Tyrosine** | 14.69491733 |
| **Cysteine** | 3.256105333 |
| **Valine** | 3.777034 |
| **Methionine** | 3.777034 |
| **Tryptophan** | 1.820329556 |
| **Phenylalanine** | 7.652614667 |
| **Isoleucine** | 2.966432 |
| **Leucine** | 4.454385778 |
| **Lysine** | 2.546316667 |
| **Proline** | 2.0 |

**Table S2.** Volatile compounds detected in the headspace of *Corydalis caseana* ssp. *brandegeei* inflorescences treated with sterile or *Metschnikowia*-fermented nectar. *RT*: Retention time, *RI*: Kovat's retention index, *Nonpolar RI*: retention index reported in the literature, *RI Ref, Column, RI type*: source and type of reported retention index, *Compound Class*: compound class, *Database Name*: Name for the top match provided by the NIST or MACE databases, *Common Name*: synonymous, common compound name, *Match Factor*: correlation score between the database mass spectrum and the pseudospectrum calculated for each detected compound, *CAS*: CAS database identifier, if applicable, *Formula*: chemical formula, *Average Peak Area:* average peak area across all floral samples, *Percent Peak Area*: percentage of the total average peak area contributed by average peak area for each compound, *Top 10 Ions:* the ten most abundant ions detected for each unidentified compound.

| RT | RI | RI non-polar | RI Ref, Column, RI type | Compound Class | Database Name | Common Name | Match Factor | CAS | Formula | Avg Peak Area | Percent Peak Area | Top 10 Ions (unknowns) |
|---|---|---|---|---|---|---|---|---|---|---|---|---|
| 0.9275 | <600 | 154 | NIST Sqalene | | Carbon dioxide | | 100 | 124389 | $CO_2$ | 388.8 | 0.01 | |
| 1.6524 | <600 | 520 | NIST OV1 | hemiterpene | Isoprene | | 97 | 78795 | $C_5H_8$ | 139406.2 | 2.34 | |
| 1.7031 | <600 | 517 | NIST DB5 | SCC | Ethanethiol | | 98 | 75081 | $C_2H_6S$ | 1806.3 | 0.03 | |
| 1.7094 | <600 | | | | Unknown 1 | | | | | 8940.0 | 0.15 | 46 (100.0%), 74 (44.4%), 59 (40.4%), 42 (13.7%), 44 (12.4%), 41 (1.4%), 75 (1.1%), 142 (1.1%), 84 (0.9%), 49 (0.8%) |
| 1.7263 | <600 | 522 | NIST DB5 VDDK | sugar catabolism | Acetic acid, methyl ester | | 98 | 79209 | $C_3H_6O_2$ | 509.4 | 0.01 | |
| 1.8223 | <600 | 553 | NIST DB1 | FAD | Cyclopentene | | 98 | 142290 | $C_5H_8$ | 20528.4 | 0.34 | |
| 1.8315 | <600 | 553 | Park et al. 2014 AT1 | SCC | Sulfurous acid, dimethyl ester | | 99 | 616422 | $C_2H_6O_3S$ | 2070.2 | 0.03 | |
| 1.9056 | <600 | | | | Unknown 68 | | | | | 636.3 | 0.01 | 46 (100.0%), 72 (73.1%), 67 (68.4%), 67 (68.4%), 41 (40.1%), 79 (26.8%), 68 (26.0%), 39 (16.8%), 42 (10.6%), 75 (8.6%) |
| 1.9212 | <600 | 568 | NIST DB5 VDDK | isoprene derivative | Methacrolein | | 99 | 78853 | $C_4H_6O$ | 3061.5 | 0.05 | |
| 1.9533 | <600 | | | | Unknown 84 | | | | | 2052.9 | 0.03 | 75 (100.0%), 60 (88.5%), 72 (30.6%), 54 (18.6%), 54 (18.6%), 42 (6.6%), 47 (4.2%), 76 (4.2%), 71 (2.9%), 52 (1.7%) |
| 2.1012 | 601 | | | | Unknown 2 | | | | | 721.4 | 0.01 | 57 (100.0%), 72 (95.8%), 42 (24.1%), 41 (22.0%), 56 (21.5%), 86 (9.7%), 86 (9.7%), 79 (8.4%), 79 (8.4%), 39 (3.9%) |
| 2.184 | 608 | 603 | NIST DB1 VDDK | hemiterpene | 3-Buten-2-ol, 2-methyl- | | 98 | 115184 | $C_5H_{10}O$ | 4149.7 | 0.07 | |
| 2.3241 | 622 | | | | Unknown 3 | | | | | 1389.7 | 0.02 | 42 (100.0%), 72 (21.4%), 41 (16.2%), 59 (16.1%), 59 (16.1%), 71 (14.0%), 39 (7.6%), 57 (7.1%), 56 (6.6%), 58 (4.1%) |
| 2.3606 | 625 | 612 | NIST HP5MS VDDK | FAD | Ethyl Acetate | | 97 | 141786 | $C_4H_8O_2$ | 2864.2 | 0.05 | |
| 2.7425 | 661 | 657 | NIST HP5 VDDK | AAD | Butanal, 3-methyl- | Isovaler-aldehyde | 98 | 590863 | $C_5H_{10}O$ | 531.4 | 0.01 | |
| 3.1392 | 699 | | | | Unknown 4 | | | | | 5486.9 | 0.09 | 44 (100.0%), 44 (100.0%), 57 (79.8%), 41 (55.9%), 71 (45.5%), 58 (32.0%), 58 (32.0%), 42 (19.7%), 56 (19.2%), 70 (19.2%) |
| 3.415 | 713 | | | | Unknown 5 | | | | | 360.2 | 0.01 | 44 (100.0%), 75 (7.2%), 59 (5.5%), 61 (5.4%), 41 (4.9%), 100 (3.8%), 46 (3.5%), 46 (3.5%), 69 (2.5%), 72 (2.4%) |
| 3.5881 | 722 | 719 | NIST DB1 | *unknown* | Cyclohexane, methyl- | | 97 | 108872 | $C_7H_{14}$ | 911.0 | 0.02 | |
| 3.7423 | 730 | | | | Unknown 6 | | | | | 347.5 | 0.01 | 41 (100.0%), 39 (14.8%), 68 (4.0%), 68 (4.0%), 73 (3.8%), 103 (3.4%), 44 (3.0%), 44 (3.0%), 38 (2.6%), 67 (2.2%) |

| | | | | | | | | | | | |
|---|---|---|---|---|---|---|---|---|---|---|---|
| 3.7732 | 731 | | | | Unknown 7 | | | | 2181.7 | 0.04 | 105 (100.0%), 77 (74.4%), 75 (40.9%), 59 (12.9%), 106 (5.9%), 106 (5.9%), 68 (5.7%), 68 (5.7%), 61 (5.0%), 76 (3.0%) |
| 3.8203 | 734 | 730 | NIST HP5MS | AAD | 1-Butanol, 3-methyl- | 98 | 123513 | C5H12O | 5599.2 | 0.09 | |
| 3.9178 | 738 | 736 | NIST HP5MS VDDK | AAD | 1-Butanol, 2-methyl- | 97 | 137326 | C5H12O | 1791.9 | 0.03 | |
| 3.9181 | 738 | 733 | NIST DB5 VDDK | isoprene derivative | Methyl Isobutyl Ketone | 93 | 108101 | C6H12O | 1722.6 | 0.03 | |
| 3.9772 | 741 | | | | Unknown 8 | | | | 378.5 | 0.01 | 84 (100.0%), 55 (30.2%), 94 (12.3%), 94 (12.3%), 83 (8.9%), 83 (8.9%), 53 (7.5%), 39 (6.1%), 44 (5.2%), 61 (4.8%) |
| 4.1315 | 749 | | | | Unknown 9 | | | | 1756.3 | 0.03 | 55 (100.0%), 58 (8.2%), 57 (4.3%), 57 (4.3%), 70 (3.3%), 79 (3.2%), 52 (3.1%), 52 (3.1%), 81 (2.9%), 42 (2.5%) |
| 4.2972 | 758 | | | | Unknown 10 | | | | 275.0 | 0.00 | 73 (100.0%), 73 (100.0%), 41 (58.0%), 88 (14.8%), 44 (9.2%), 44 (9.2%), 42 (7.1%), 137 (6.4%), 137 (6.4%), 57 (5.2%) |
| 4.5088 | 768 | | | | Unknown 11 | | | | 301.4 | 0.01 | 42 (100.0%), 73 (42.6%), 73 (42.6%), 59 (31.4%), 59 (31.4%), 74 (15.9%), 69 (10.5%), 170 (8.2%), 170 (8.2%), 44 (7.5%) |
| 4.5964 | 773 | 764 | NIST DB5MS | AAD | Isobutyl acetate | 96 | 110190 | C6H12O2 | 677.7 | 0.01 | |
| 4.6853 | 777 | 780 | NIST HP5MS VDDK | FAD | Butanoic acid, 2-methyl-, methyl ester | 99 | 868575 | C6H12O2 | 25356.3 | 0.43 | |
| 4.692 | 777 | | | | Unknown 12 | | | | 2306.5 | 0.04 | 88 (100.0%), 88 (100.0%), 57 (15.6%), 57 (15.6%), 41 (8.0%), 74 (5.1%), 101 (4.6%), 59 (4.4%), 56 (3.9%), 85 (3.9%) |
| 4.7208 | 779 | 765 | NIST HP5MS | FAD | Methyl isovalerate | 98 | 556241 | C6H12O2 | 22435.7 | 0.38 | |
| 4.9358 | 790 | | | | Unknown 13 | | | | 25342.7 | 0.42 | 55 (100.0%), 56 (45.2%), 41 (42.6%), 42 (25.4%), 84 (23.9%), 70 (23.3%), 39 (15.6%), 69 (12.2%), 83 (11.4%), 57 (5.7%) |
| 4.9451 | 790 | 794 | NIST DB5 VDDK | FAD | Cyclopentanone | 98 | 120923 | C5H8O | 61958.7 | 1.04 | |
| 4.9725 | 792 | | | | Unknown 14 | | | | 22416.6 | 0.38 | 56 (100.0%), 41 (89.3%), 70 (79.0%), 70 (79.0%), 42 (68.3%), 69 (40.4%), 83 (35.5%), 39 (28.3%), 57 (14.6%), 82 (9.8%) |
| 5.1056 | 798 | | | | Unknown 15 | | | | 965.6 | 0.02 | 86 (100.0%), 39 (20.0%), 41 (11.0%), 133 (6.7%), 149 (5.2%), 59 (4.8%), 44 (3.8%), 44 (3.8%), 73 (2.9%), 38 (2.8%) |
| 5.1527 | 800 | | | | Unknown 16 | | | | 4533.2 | 0.08 | 85 (100.0%), 71 (44.6%), 70 (28.5%), 57 (22.8%), 84 (20.6%), 84 (20.6%), 114 (16.1%), 41 (10.8%), 42 (10.6%), 86 (5.7%) |
| 5.1795 | 801 | 800 | by definition | *likely contaminant* | Octane | 95 | 111659 | C8H18 | 16546.0 | 0.28 | |
| 5.2867 | 805 | | | | Unknown 17 | | | | 2646.1 | 0.04 | 46 (100.0%), 44 (3.3%), 44 (3.3%), 100 (2.2%), 47 (1.7%), 56 (1.3%), 59 (1.2%), 67 (1.2%), 67 (1.2%), 166 (1.2%) |

| | | | | | | | | | | | | |
|---|---|---|---|---|---|---|---|---|---|---|---|---|
| 5.4071 | 810 | | | | Unknown 69 | | | | | 470.6 | 0.01 | 57 (100.0%), 166 (42.0%), 164 (34.0%), 129 (31.5%), 131 (27.8%), 168 (17.9%), 94 (11.7%), 41 (8.5%), 133 (7.5%), 98 (4.5%) |
| 5.5054 | 813 | | | | Unknown 18 | | | | | 4694.7 | 0.08 | 60 (100.0%), 42 (10.2%), 44 (3.0%), 61 (2.2%), 61 (2.2%), 77 (1.7%), 77 (1.7%), 46 (1.3%), 46 (1.3%), 55 (1.3%) |
| 5.5521 | 815 | 820 | NIST OV101 | *likely contaminant* | N,N-Dimethylacetamide | | 93 | 127195 | C4H9NO | 734.5 | 0.01 | |
| 5.6074 | 817 | 808 | NIST DB5 Norm | *likely contaminant* | Isobutyl ether | | 97 | 628557 | C8H18O | 616.8 | 0.01 | |
| 5.82 | 824 | | | | Unknown 70 | | | | | 352.3 | 0.01 | 207 (100.0%), 67 (43.0%), 54 (26.1%), 54 (26.1%), 94 (24.7%), 94 (24.7%), 109 (19.5%), 110 (16.9%), 110 (16.9%), 72 (16.1%) |
| 5.878 | 826 | | | | Unknown 71 | | | | | 913.1 | 0.02 | 207 (100.0%), 207 (100.0%), 70 (52.8%), 42 (48.0%), 55 (43.1%), 208 (21.5%), 41 (10.0%), 94 (7.8%), 209 (7.4%), 74 (6.1%) |
| 6.0659 | 833 | | | | Unknown 19 | | | | | 7585.9 | 0.13 | 77 (100.0%), 78 (3.5%), 57 (2.1%), 57 (2.1%), 44 (1.9%), 96 (1.8%), 96 (1.8%), 79 (1.7%), 79 (1.7%), 72 (1.6%) |
| 6.104 | 835 | 835 | NIST DB5 VDDK | FAD | 2-Cyclopenten-1-one | | 94 | 930303 | C5H6O | 458.1 | 0.01 | |
| 6.2391 | 839 | 836 | NIST DB5 | FAD | Cyclopentanone, 2-methyl- | | 98 | 1120725 | C6H10O | 3276.5 | 0.05 | |
| 6.265 | 840 | 827 | NIST DB5 VDDK | FAD | Formic acid, pentyl ester | pentyl formate | 92 | 638493 | C6H12O2 | 1176.7 | 0.02 | |
| 6.3264 | 843 | 839 | NIST HP5MS Norm | *unknown* | 2-Pentanone, 4-hydroxy-4-methyl- | diacetone alcohol | 94 | 123422 | C6H12O2 | 594.8 | 0.01 | |
| 6.7156 | 856 | 855 | NIST DB5MS | FAD | 3-Hexen-1-ol, (E)- | | 98 | 928972 | C6H12O | 7043.8 | 0.12 | |
| 7.2638 | 876 | 876 | NIST DB5 VDDK | benzenoid | Phenylethyne | | 99 | 536743 | C8H6 | 430.0 | 0.01 | |
| 7.3671 | 880 | 880 | NIST HP5MS VDDK | AAD | 1-Butanol, 2-methyl-, acetate | | 97 | 624419 | C7H14O2 | 1099.1 | 0.02 | |
| 7.7325 | 893 | | | | Unknown 72 | | | | | 525.8 | 0.01 | 91 (100.0%), 55 (44.0%), 42 (30.3%), 106 (29.1%), 98 (29.0%), 98 (29.0%), 68 (11.8%), 67 (9.7%), 73 (9.4%), 105 (8.2%) |
| 7.7839 | 895 | 897 | NIST HP5MS VDDK | FAD | 5-Hexenal, 4-methylene- | | 99 | 17844212 | C7H10O | 534.1 | 0.01 | |
| 7.9905 | 902 | | | | Unknown 20 | | | | | 2255.2 | 0.04 | 70 (100.0%), 44 (50.1%), 42 (19.0%), 81 (10.9%), 55 (10.2%), 55 (10.2%), 68 (9.5%), 68 (9.5%), 86 (9.2%), 41 (6.5%) |
| 8.1709 | 908 | 909 | NIST DB5 VDDK | *likely contaminant* | Ethanol, 2-butoxy- | | 100 | 111762 | C6H14O2 | 14043.7 | 0.24 | |
| 8.3473 | 914 | | | | Unknown 21 | | | | | 546.1 | 0.01 | 42 (100.0%), 86 (19.3%), 41 (15.4%), 56 (11.5%), 85 (5.5%), 69 (4.4%), 93 (4.3%), 98 (3.4%), 39 (2.4%), 57 (1.7%) |

| | | | | | | | | | | | | |
|---|---|---|---|---|---|---|---|---|---|---|---|---|
| 8.5132 | 919 | 921 | NIST HP5MS VDDK | monoterpene | 5,5-Dimethyl-1-vinylbicyclo[2.1.1]hexane | | 98 | 16626394 | C10H16 | 13310.1 | 0.22 | |
| 8.5767 | 921 | | | | Unknown 73 | | | | | 1492.1 | 0.03 | 54 (100.0%), 66 (49.4%), 42 (41.7%), 71 (17.1%), 93 (15.3%), 93 (15.3%), 39 (11.3%), 41 (8.2%), 67 (8.2%), 67 (8.2%) |
| 8.7831 | 928 | 925 | NIST HP5MS | monoterpene | Bicyclo[3.1.0]hex-2-ene, 2-methyl-5-(1-methylethyl)- | alpha thujene | 99 | 2867052 | C10H16 | 8601.4 | 0.14 | |
| 8.8642 | 931 | 933 | NIST DB5 VDDK | FAD | Ethanone, 1-cyclopentyl- | | 89 | 6004600 | C7H12O | 421.8 | 0.01 | |
| 8.985 | 935 | 935 | NIST DB5 | monoterpene | (1R)-2,6,6-Trimethylbicyclo[3.1.1]hept-2-ene | alpha pinene (dextro) | 99 | 7785708 | C10H16 | 143293.1 | 2.40 | |
| 9.1154 | 939 | | | | Unknown 22 | | | | | 2114.4 | 0.04 | 84 (100.0%), 84 (100.0%), 112 (32.5%), 56 (32.4%), 83 (26.4%), 68 (9.1%), 42 (8.5%), 55 (7.1%), 55 (7.1%), 41 (6.9%) |
| 9.1186 | 939 | 941 | Schäfer et al 2025 ZB5HT | FAD | Cyclopentanone, 2-ethyl- | | 97 | 4971180 | C7H12O | 11474.2 | 0.19 | |
| 9.1241 | 940 | 944 | NIST SE30 | NCC | 1-Butylpyrrolidine | | 98 | 767102 | C8H17N | 1429.1 | 0.02 | |
| 10.0495 | 970 | 972 | NIST HP5MS Norm | monoterpene | Bicyclo[3.1.1]hept-2-ene, 3,6,6-trimethyl- | pinane | 96 | 4889832 | C10H16 | 419.8 | 0.01 | |
| 10.2113 | 976 | 978 | NIST HP5 | monoterpene | Bicyclo[3.1.0]hex-2-ene, 4-methyl-1-(1-methylethyl)- | beta thujene | 99 | 28634891 | C10H16 | 192246.9 | 3.22 | |
| 10.4139 | 982 | | | | Unknown 23 | | | | | 950.8 | 0.02 | 57 (100.0%), 87 (63.8%), 69 (56.7%), 69 (56.7%), 41 (46.6%), 56 (46.1%), 118 (12.8%), 72 (10.9%), 103 (10.5%), 117 (9.1%) |
| 10.4657 | 984 | 988 | NIST DB5 VDDK | benzenoid | .alpha.-Methylstyrene | | 84 | 98839 | C9H10 | 766.6 | 0.01 | |
| 10.5089 | 985 | | | | Unknown 24 | | | | | 1807.3 | 0.03 | 207 (100.0%), 223 (18.0%), 103 (17.4%), 208 (15.4%), 209 (9.9%), 191 (8.1%), 76 (4.8%), 133 (4.2%), 133 (4.2%), 104 (4.0%) |
| 10.745 | 993 | 991 | NIST DB5 | monoterpene | .beta.-Myrcene | | 99 | 123353 | C10H16 | 1768086.9 | 29.64 | |
| 10.8233 | 996 | | | | Unknown 25 | | | | | 5705.5 | 0.10 | 90 (100.0%), 87 (49.2%), 41 (14.9%), 69 (9.9%), 93 (8.0%), 57 (7.9%), 58 (5.3%), 75 (3.8%), 62 (2.7%), 91 (2.5%) |
| 10.9821 | 1001 | | | | Unknown 26 | | | | | 16342.3 | 0.27 | 281 (100.0%), 281 (100.0%), 57 (24.0%), 282 (22.6%), 283 (13.0%), 265 (12.9%), 84 (9.7%), 56 (9.5%), 44 (9.2%), 41 (8.6%) |
| 11.0856 | 1006 | 1007 | NIST HP5MS | monoterpene | .alpha.-Phellandrene | | 94 | 99832 | C10H16 | 1168.7 | 0.02 | |
| 11.1633 | 1009 | 1013 | NIST VF5MS VDDK | FAD | 4-Hexen-1-ol, (4E)-, acetate | | 98 | 1000352719 | C8H14O2 | 7136.2 | 0.12 | |

| | | | | | | | | | | | | |
|---|---|---|---|---|---|---|---|---|---|---|---|---|
| 11.3089 | 1015 | | | monoterpene | Unknown monoterpene 1 | | | | | 4200.7 | 0.07 | 93 (100.0%), 69 (60.0%), 41 (57.2%), 91 (12.3%), 79 (9.3%), 77 (7.9%), 94 (6.9%), 80 (6.8%), 39 (5.4%), 53 (5.1%) |
| 11.3283 | 1016 | | | | Unknown 27 | | | | | 1399.9 | 0.02 | 69 (100.0%), 56 (88.0%), 56 (88.0%), 41 (61.7%), 84 (36.7%), 61 (36.3%), 55 (31.4%), 93 (30.0%), 93 (30.0%), 42 (24.9%) |
| 11.3392 | 1017 | 1014 | NIST HP5 VDDK | FAD | Acetic acid, hexyl ester | hexyl acetate | 94 | 142927 | C8H16O2 | 969.4 | 0.02 | |
| 11.408 | 1019 | 1017 | NIST HP5MS | monoterpene | 1,3-Cyclohexadiene, 1-methyl-4-(1-methylethyl)- | alpha terpinene | 96 | 99865 | C10H16 | 547.6 | 0.01 | |
| 11.4727 | 1022 | | | monoterpene | Unknown monoterpene 2 | | | | | 819.2 | 0.01 | 79 (100.0%), 91 (56.8%), 119 (32.1%), 77 (21.3%), 77 (21.3%), 92 (20.7%), 78 (15.5%), 105 (10.2%), 105 (10.2%), 53 (9.1%) |
| 11.6092 | 1028 | 1030 | NIST DB5 | benzenoid | Benzene, 1-methyl-3-(1-methylethyl)- | m-Cymene | 95 | 535773 | C10H14 | 1751.0 | 0.03 | |
| 11.7252 | 1033 | 1031 | NIST DB5 | monoterpene | D-Limonene | | 99 | 5989275 | C10H16 | 950110.9 | 15.93 | |
| 11.9434 | 1042 | 1041 | NIST HP5MS | monoterpene | trans-.beta.-Ocimene | | 98 | 3779611 | C10H16 | 138149.9 | 2.32 | |
| 12.0691 | 1048 | | | | Unknown 28 | | | | | 893.0 | 0.01 | 91 (100.0%), 91 (100.0%), 120 (14.3%), 65 (12.1%), 87 (6.9%), 92 (6.7%), 63 (4.6%), 63 (4.6%), 56 (4.0%), 116 (4.0%) |
| 12.1513 | 1051 | 1044 | NIST HP5MS | monoterpene | Limonene | | 97 | 138863 | C10H16 | 5264.1 | 0.09 | |
| 12.1855 | 1053 | 1050 | NIST HP5 | monoterpene | .beta.-Ocimene | | 97 | 13877913 | C10H16 | 25806.4 | 0.43 | |
| 12.5996 | 1070 | 1065 | NIST HP5MS | benzenoid | Acetophenone | | 97 | 98862 | C8H8O | 2671.0 | 0.04 | |
| 12.7261 | 1076 | | | | Unknown 29 | | | | | 1845.7 | 0.03 | 59 (100.0%), 71 (10.9%), 58 (10.6%), 99 (10.6%), 55 (7.2%), 123 (6.7%), 123 (6.7%), 56 (5.6%), 85 (5.3%), 67 (5.2%) |
| 12.737 | 1076 | | | | Unknown 30 | | | | | 515.1 | 0.01 | 59 (100.0%), 58 (44.6%), 71 (40.9%), 71 (40.9%), 99 (40.2%), 85 (31.9%), 57 (20.9%), 57 (20.9%), 105 (10.3%), 105 (10.3%) |
| 12.9889 | 1087 | 1086 | NIST HP5MS VDDK | sugar catabolism | Furyl hydroxymethyl ketone | | 97 | 17678192 | C6H6O3 | 354.8 | 0.01 | |
| 12.9906 | 1087 | | | | Unknown 31 | | | | | 7673.9 | 0.13 | 126 (100.0%), 55 (18.5%), 95 (7.5%), 70 (4.0%), 83 (3.6%), 85 (3.6%), 85 (3.6%), 127 (2.7%), 69 (2.0%), 56 (1.9%) |
| 13.0227 | 1088 | | | | Unknown 32 | | | | | 2310.3 | 0.04 | 207 (100.0%), 208 (18.2%), 209 (16.6%), 193 (9.3%), 121 (5.8%), 191 (4.7%), 105 (3.3%), 60 (2.9%), 60 (2.9%), 97 (2.5%) |
| 13.143 | 1093 | | | | Unknown 33 | | | | | 258.9 | 0.00 | 58 (100.0%), 58 (100.0%), 56 (42.2%), 70 (28.4%), 83 (22.2%), 83 (22.2%), 69 (19.0%), 55 (18.4%), 71 (17.9%), 84 (17.0%) |
| 13.1868 | 1095 | | | | Unknown 34 | | | | | 799.4 | 0.01 | 79 (100.0%), 71 (42.3%), 59 (29.4%), 41 (23.9%), 67 (16.0%), 81 (12.7%), 39 (10.7%), 85 (9.9%), 85 (9.9%), 53 (7.8%) |

| | | | | | | | | | | | | |
|---|---|---|---|---|---|---|---|---|---|---|---|---|
| 13.2335 | 1097 | 1088 | Meng et al. 2024 DB5-MS | *likely contaminant* | Benzoic acid, hydrazide | | 100 | 613945 | C7H8N2O | 63985.2 | 1.07 | |
| 13.3593 | 1103 | | | | Unknown 85 | | | | | 7294.2 | 0.12 | 119 (100.0%), 59 (92.0%), 79 (87.5%), 91 (69.8%), 81 (66.2%), 93 (56.2%), 67 (52.0%), 80 (51.1%), 95 (40.7%), 94 (37.3%) |
| 13.4758 | 1109 | 1106 | NIST DB5 | benzenoid | Benzoic acid, methyl ester | methyl benzoate | 99 | 93583 | C8H8O2 | 444.1 | 0.01 | |
| 13.6433 | 1118 | 1121 | NIST DB5 | benzenoid | Phenylethyl Alcohol | | 100 | 60128 | C8H10O | 1074339.9 | 18.01 | |
| 13.7717 | 1125 | | | | Unknown 35 | | | | | 2181.7 | 0.04 | 91 (100.0%), 119 (17.9%), 107 (16.8%), 59 (15.9%), 79 (12.7%), 92 (11.6%), 92 (11.6%), 135 (11.5%), 134 (9.4%), 65 (8.3%) |
| 13.8183 | 1128 | | | | Unknown 74 | | | | | 1202.1 | 0.02 | 93 (100.0%), 77 (66.8%), 78 (27.4%), 58 (16.8%), 53 (15.2%), 53 (15.2%), 94 (13.2%), 94 (13.2%), 55 (11.4%), 98 (7.0%) |
| 13.8244 | 1128 | | | | Unknown 36 | | | | | 3153.4 | 0.05 | 91 (100.0%), 93 (78.3%), 77 (63.8%), 77 (63.8%), 92 (31.3%), 78 (20.9%), 58 (18.2%), 53 (17.7%), 65 (12.0%), 65 (12.0%) |
| 13.8886 | 1131 | 1125 | NIST SE30 | benzenoid | Benzeneethanamine | | 90 | 64040 | C8H11N | 5918.9 | 0.10 | |
| 13.9125 | 1133 | 1131 | NIST HP5MS VDDK | monoterpene | 2,4,6-Octatriene, 2,6-dimethyl-, (E,Z)- | allo ocimene | 99 | 7216560 | C10H16 | 13082.0 | 0.22 | |
| 14.1055 | 1143 | 1143 | NIST HP5MS | benzenoid | Benzyl nitrile | | 99 | 140294 | C8H7N | 43230.9 | 0.72 | |
| 14.1309 | 1144 | 1132 | NIST MethylSil | monoterpene | 1,7-Octadien-3-one, 2-methyl-6-methylene- | | 98 | 41702607 | C10H14O | 40223.1 | 0.67 | |
| 14.2267 | 1150 | 1154 | NIST HP5MS VDDK | sugar catabolism | 4H-Pyran-4-one, 2,3-dihydro-3,5-dihydroxy-6-methyl- | Hydroxy-dihydro-maltol | 98 | 28564832 | C6H8O4 | 1198.9 | 0.02 | |
| 14.2655 | 1152 | 1148 | NIST HP5 | monoterpene | Camphor | | 95 | 76222 | C10H16O | 528.6 | 0.01 | |
| 14.3105 | 1154 | 1159 | NIST 5PMS VDDK | FAD | Acetic acid, 2-ethylhexyl ester | | 98 | 103093 | C10H20O2 | 7250.5 | 0.12 | |
| 14.4417 | 1161 | | | monoterpene | Unknown monoterpene 3 | | | | | 4860.5 | 0.08 | 67 (100.0%), 69 (78.2%), 41 (74.0%), 71 (62.0%), 68 (59.4%), 79 (56.2%), 84 (43.1%), 105 (30.6%), 39 (29.3%), 93 (29.3%) |
| 14.4865 | 1163 | | | | Unknown 37 | | | | | 204.6 | 0.00 | 73 (100.0%), 267 (43.0%), 70 (25.8%), 55 (18.6%), 83 (18.6%), 83 (18.6%), 41 (13.2%), 57 (11.0%), 56 (9.0%), 268 (8.4%) |
| 14.5219 | 1165 | | | | Unknown 38 | | | | | 1693.1 | 0.03 | 79 (100.0%), 80 (57.5%), 175 (18.9%), 244 (9.9%), 244 (9.9%), 125 (7.4%), 81 (6.4%), 110 (4.8%), 110 (4.8%), 68 (4.7%) |
| 14.5688 | 1168 | 1170 | NIST HP5 VDDK | benzenoid | Benzene, 1,4-dimethoxy- | | 99 | 150787 | C8H10O2 | 28575.1 | 0.48 | |
| 14.578 | 1168 | 1165 | NIST HP5MS VDDK | benzenoid | Acetic acid, phenylmethyl ester | benzyl acetate | 98 | 140114 | C9H10O2 | 1393.1 | 0.02 | |

| | | | | | | | | | | | | |
|---|---|---|---|---|---|---|---|---|---|---|---|---|
| 14.579 | 1168 | | | | Unknown 39 | | | | | 369.8 | 0.01 | 108 (100.0%), 53 (47.1%), 107 (31.3%), 81 (30.3%), 81 (30.3%), 79 (26.2%), 125 (23.9%), 135 (21.7%), 105 (19.1%), 54 (15.4%) |
| 14.7067 | 1175 | 1176 | NIST HP5MS VDDK | FAD | Octanoic acid | | 82 | 124072 | C8H16O2 | 1481.6 | 0.02 | |
| 14.7672 | 1178 | | | | Unknown 40 | | | | | 949.5 | 0.02 | 134 (100.0%), 91 (92.3%), 93 (63.9%), 106 (35.3%), 92 (34.0%), 119 (33.4%), 119 (33.4%), 66 (16.9%), 105 (16.6%), 78 (13.0%) |
| 14.8031 | 1180 | | | | Unknown 41 | | | | | 1490.1 | 0.02 | 104 (100.0%), 117 (18.3%), 117 (18.3%), 90 (11.3%), 91 (9.3%), 109 (9.2%), 103 (5.3%), 78 (4.2%), 65 (3.9%), 118 (3.5%) |
| 14.8104 | 1181 | | | | Unknown 42 | | | | | 2293.5 | 0.04 | 117 (100.0%), 117 (100.0%), 90 (57.2%), 91 (46.6%), 91 (46.6%), 104 (22.6%), 118 (11.1%), 149 (10.5%), 65 (10.3%), 89 (9.4%) |
| 14.8297 | 1182 | 1187 | NIST HP5MS | benzenoid | Benzeneacetic acid, methyl ester | | 99 | 101417 | C9H10O2 | 6569.4 | 0.11 | |
| 14.8495 | 1183 | 1174 | NIST SE54 Norm | monoterpene | p-Mentha-1,8-dien-4-ol | Limonen-4-ol | 91 | | C10H16O | 792.0 | 0.01 | |
| 14.8605 | 1183 | 1178 | NIST HP5MS | monoterpene | Terpinen-4-ol | | 97 | 562743 | C10H18O | 226.0 | 0.00 | |
| 15.0988 | 1196 | 1191 | NIST HP5MS | monoterpene | .alpha.-Terpineol | | 99 | 98555 | C10H18O | 14276.0 | 0.24 | |
| 15.3675 | 1212 | | | | Unknown 75 | | | | | 1175.3 | 0.02 | 55 (100.0%), 68 (56.7%), 72 (46.1%), 72 (46.1%), 41 (40.9%), 59 (31.7%), 67 (30.6%), 67 (30.6%), 96 (28.9%), 69 (28.1%) |
| 15.4331 | 1216 | | | | Unknown 43 | | | | | 870.1 | 0.01 | 57 (100.0%), 71 (93.5%), 71 (93.5%), 85 (54.8%), 97 (42.2%), 70 (37.3%), 55 (33.4%), 56 (33.0%), 56 (33.0%), 41 (31.1%) |
| 15.5739 | 1225 | 1220 | NIST HP5MS | monoterpene | 2-Cyclohexen-1-ol, 2-methyl-5-(1-methylethenyl)-, cis- | carveol | 89 | 1197064 | C10H16O | 507.0 | 0.01 | |
| 15.6256 | 1228 | | | | Unknown 44 | | | | | 208.1 | 0.00 | 119 (100.0%), 59 (35.7%), 134 (31.3%), 134 (31.3%), 79 (18.7%), 91 (17.5%), 120 (10.5%), 120 (10.5%), 95 (7.7%), 67 (6.6%) |
| 15.7371 | 1235 | 1233 | NIST HP5MS VDDK | sugar dehydration | 5-Hydroxymethylfurfural | | 98 | 67470 | C6H6O3 | 6238.6 | 0.10 | |
| 15.9255 | 1247 | | | | Unknown 45 | | | | | 386.6 | 0.01 | 84 (100.0%), 67 (24.5%), 93 (21.6%), 108 (18.5%), 55 (18.3%), 83 (18.2%), 103 (17.3%), 41 (14.9%), 69 (12.6%), 98 (11.1%) |
| 15.9417 | 1248 | | | | Unknown 46 | | | | | 457.8 | 0.01 | 105 (100.0%), 105 (100.0%), 77 (45.6%), 125 (14.5%), 84 (13.1%), 84 (13.1%), 98 (12.4%), 71 (10.1%), 70 (9.4%), 51 (8.5%) |
| 15.995 | 1252 | 1249 | NIST DB5 VDDK | monoterpene | D-Carvone | | 98 | 2244168 | C10H14O | 1457.6 | 0.02 | |
| 15.9973 | 1252 | 1230 | PubChem | FAD | Glycerol 1,2-diacetate | diacetin | 93 | 102625 | C7H12O5 | 960.1 | 0.02 | |

| | | | | | | | | | | | |
|---|---|---|---|---|---|---|---|---|---|---|---|
| 16.1075 | 1259 | | | | Unknown 47 | | | | 5894.2 | 0.10 | 91 (100.0%), 117 (91.4%), 90 (47.3%), 118 (34.9%), 135 (27.1%), 89 (16.9%), 65 (15.8%), 119 (11.3%), 92 (9.8%), 116 (7.0%) |
| 16.1664 | 1262 | 1256 | NIST HP5MS | benzenoid | Acetic acid, 2-phenylethyl ester | 99 | 103457 | C10H12O2 | 313174.6 | 5.25 | |
| 16.226 | 1266 | | | | Unknown 48 | | | | 13241.2 | 0.22 | 104 (100.0%), 113 (18.1%), 56 (7.2%), 105 (6.0%), 78 (4.8%), 78 (4.8%), 91 (4.7%), 84 (4.1%), 85 (4.0%), 85 (4.0%) |
| 16.4467 | 1280 | | | | Unknown 49 | | | | 11587.4 | 0.19 | 117 (100.0%), 91 (86.5%), 90 (56.8%), 90 (56.8%), 118 (26.2%), 118 (26.2%), 89 (20.9%), 135 (19.3%), 65 (19.2%), 92 (12.1%) |
| 16.5012 | 1283 | 1289 | NIST HP5 VDDK | | Indole | 94 | 120729 | C8H7N | 715.3 | 0.01 | |
| 16.6661 | 1294 | 1282 | Cui et al. 2017 - HP5 | FAD (Weber 2002) | [1,1'-Bicyclopentyl]-2-one | 100 | 4884246 | C10H16O | 323400.7 | 5.42 | |
| 16.7273 | 1297 | | | | Unknown 50 | | | | 14878.7 | 0.25 | 84 (100.0%), 67 (12.0%), 83 (7.5%), 83 (7.5%), 54 (2.3%), 81 (2.1%), 152 (1.8%), 135 (1.6%), 58 (1.5%), 68 (1.3%) |
| 16.7937 | 1302 | 1292 | NIST DB5 | NCC | Indole | 100 | 120729 | C8H7N | 105742.7 | 1.77 | |
| 16.8507 | 1306 | 1306 | NIST HP5MS VDDK | benzenoid | Benzene, (2-nitroethyl)- | 98 | 6125242 | C8H9NO2 | 15461.9 | 0.26 | |
| 16.9644 | 1314 | | | | Unknown 51 | | | | 534.2 | 0.01 | 281 (100.0%), 282 (24.9%), 283 (15.7%), 283 (15.7%), 72 (9.0%), 72 (9.0%), 265 (8.2%), 265 (8.2%), 249 (6.3%), 114 (5.7%) |
| 17.088 | 1322 | | | | Unknown 76 | | | | 339.8 | 0.01 | 281 (100.0%), 267 (71.0%), 73 (29.9%), 282 (23.2%), 268 (20.2%), 81 (14.5%), 269 (14.1%), 283 (14.0%), 141 (7.6%), 150 (6.7%) |
| 17.0885 | 1322 | | | | Unknown 77 | | | | 635.2 | 0.01 | 73 (100.0%), 281 (88.4%), 267 (81.0%), 282 (26.0%), 268 (17.8%), 81 (17.3%), 269 (14.0%), 283 (14.0%), 82 (11.3%), 141 (10.9%) |
| 17.0963 | 1323 | 1318 | NIST HP5MS VDDK | monoterpene | 6-Hydroxycarvotan acetone | 93 | 131486992 | C10H16O2 | 1246.1 | 0.02 | |
| 17.165 | 1328 | | | | Unknown 78 | | | | 3655.3 | 0.06 | 147 (100.0%), 74 (46.6%), 207 (27.2%), 207 (27.2%), 75 (21.4%), 148 (17.4%), 251 (12.1%), 267 (11.4%), 249 (8.4%), 249 (8.4%) |
| 17.2481 | 1334 | | | | Unknown 52 | | | | 403.7 | 0.01 | 75 (100.0%), 105 (76.2%), 104 (24.7%), 104 (24.7%), 129 (23.0%), 60 (14.0%), 98 (8.9%), 87 (8.5%), 77 (8.3%), 77 (8.3%) |
| 17.2743 | 1336 | | | | Unknown 53 | | | | 1561.7 | 0.03 | 98 (100.0%), 67 (21.8%), 67 (21.8%), 41 (12.4%), 69 (10.6%), 87 (9.7%), 87 (9.7%), 59 (7.0%), 55 (6.4%), 95 (5.5%) |
| 17.3231 | 1339 | | | | Unknown 79 | | | | 376.2 | 0.01 | 98 (100.0%), 131 (42.1%), 57 (30.1%), 118 (23.9%), 91 (23.2%), 160 (22.6%), 145 |

| | | | | | | | | | | | | |
|---|---|---|---|---|---|---|---|---|---|---|---|---|
| | | | | | | | | | | | | (22.2%), 67 (18.8%), 117 (17.2%), 105 (13.1%) |
| 17.5271 | 1353 | | | | Unknown 80 | | | | | 1220.1 | 0.02 | 79 (100.0%), 150 (96.3%), 122 (63.7%), 108 (62.3%), 93 (58.5%), 93 (58.5%), 109 (52.6%), 109 (52.6%), 135 (34.9%), 107 (30.6%) |
| 17.5272 | 1353 | 1347 | NIST HP5 VDDK | monoterpene | 2-Cyclohexen-1-one, 3-methyl-6-(1-methylethylidene)- | Piperiten-one | 88 | 491098 | C10H14O | 1617.9 | 0.03 | |
| 17.9904 | 1386 | | | | Unknown 54 | | | | | 342.7 | 0.01 | 70 (100.0%), 71 (34.9%), 71 (34.9%), 55 (33.6%), 83 (18.7%), 57 (15.3%), 139 (7.3%), 41 (7.1%), 84 (4.4%), 84 (4.4%) |
| 18.1692 | 1399 | | | | Unknown 81 | | | | | 1119.7 | 0.02 | 119 (100.0%), 93 (74.8%), 58 (60.0%), 91 (30.4%), 150 (27.4%), 79 (25.6%), 79 (25.6%), 59 (17.4%), 77 (15.5%), 105 (13.4%) |
| 18.2663 | 1406 | | | | Unknown 55 | | | | | 623.8 | 0.01 | 73 (100.0%), 142 (10.8%), 59 (9.1%), 129 (5.6%), 74 (5.4%), 75 (5.1%), 143 (3.6%), 135 (2.9%), 147 (2.6%), 117 (1.9%) |
| 18.6816 | 1437 | | | | Unknown 56 | | | | | 705.5 | 0.01 | 112 (100.0%), 84 (96.5%), 67 (40.3%), 83 (20.4%), 41 (18.8%), 55 (14.0%), 124 (9.9%), 113 (8.0%), 95 (7.8%), 85 (7.0%) |
| 18.9206 | 1455 | | | | Unknown 57 | | | | | 477.3 | 0.01 | 142 (100.0%), 100 (80.7%), 57 (16.5%), 44 (12.3%), 58 (12.2%), 58 (12.2%), 86 (8.9%), 86 (8.9%), 158 (8.1%), 104 (7.6%) |
| 18.9495 | 1458 | 1451 | Bhattacharya et al. 2024 DB5 | NCC | Quinoline, 1,2-dihydro-2,2,4-trimethyl- | | 97 | 147477 | C12H15N | 600.1 | 0.01 | |
| 19.0102 | 1462 | 1460 | NIST HP5MS | sesquiterpene | (E)-.beta.-Famesene | | 98 | 18794848 | C15H24 | 5662.1 | 0.09 | |
| 19.3532 | 1488 | | | | Unknown 58 | | | | | 1396.2 | 0.02 | 73 (100.0%), 267 (17.4%), 104 (11.6%), 223 (11.5%), 147 (9.2%), 281 (7.7%), 221 (6.4%), 74 (5.9%), 207 (4.1%), 268 (3.1%) |
| 19.5242 | 1501 | | | | Unknown 59 | | | | | 3132.8 | 0.05 | 57 (100.0%), 85 (21.2%), 71 (11.9%), 41 (8.5%), 70 (5.9%), 70 (5.9%), 55 (5.2%), 119 (3.8%), 56 (3.7%), 42 (3.6%) |
| 19.58 | 1506 | | | | Unknown 60 | | | | | 204.5 | 0.00 | 158 (100.0%), 159 (56.7%), 130 (13.5%), 130 (13.5%), 131 (10.7%), 119 (9.7%), 149 (7.4%), 93 (6.8%), 160 (5.1%), 99 (3.0%) |
| 19.6733 | 1513 | 1509 | NIST HP5 | sesquiterpene | .alpha.-Farnesene | | 94 | 502614 | C15H24 | 7656.1 | 0.13 | |
| 20.3171 | 1566 | | | | Unknown 82 | | | | | 225.6 | 0.00 | 153 (100.0%), 153 (100.0%), 57 (67.0%), 57 (67.0%), 71 (41.3%), 85 (34.2%), 73 (26.2%), 73 (26.2%), 196 (18.2%), 181 (17.0%) |
| 21.0855 | 1631 | | | | Unknown 61 | | | | | 389.1 | 0.01 | 58 (100.0%), 72 (50.2%), 100 (50.2%), 101 (31.4%), 101 (31.4%), 98 (31.0%), 198 (26.8%), 44 (19.4%), 116 (10.6%), 116 (10.6%) |

| | | | | | | | | | | | |
|---|---|---|---|---|---|---|---|---|---|---|---|
| 21.2936 | 1649 | | | | Unknown 62 | | | | 421.1 | 0.01 | 105 (100.0%), 105 (100.0%), 109 (44.9%), 77 (42.2%), 213 (26.2%), 213 (26.2%), 151 (22.7%), 151 (22.7%), 182 (20.2%), 51 (12.2%) |
| 21.3531 | 1654 | 1643 | NIST HP5MS Norm | *likely contaminant* | Benzene, (1-propyloctyl)- | 92 | 4536861 | C17H28 | 422.9 | 0.01 | |
| 21.4825 | 1665 | | | | Unknown 63 | | | | 8726.1 | 0.15 | 57 (100.0%), 71 (56.2%), 85 (33.0%), 85 (33.0%), 41 (13.8%), 119 (11.5%), 99 (10.5%), 55 (9.0%), 55 (9.0%), 113 (7.4%) |
| 21.672 | 1682 | 1669 | NIST DB1 | FAD | 1-Tetradecanol | 97 | 112721 | C14H30O | 904.4 | 0.02 | |
| 21.7783 | 1691 | | | | Unknown 64 | | | | 138.0 | 0.00 | 85 (100.0%), 120 (12.1%), 120 (12.1%), 100 (10.1%), 128 (7.4%), 137 (7.3%), 137 (7.3%), 81 (4.3%), 119 (3.7%), 121 (3.7%) |
| 21.831 | 1696 | | | | Unknown 65 | | | | 1779.7 | 0.03 | 83 (100.0%), 83 (100.0%), 69 (90.9%), 69 (90.9%), 97 (86.4%), 70 (70.8%), 57 (66.5%), 111 (50.9%), 56 (47.5%), 84 (45.7%) |
| 22.0265 | 1713 | 1715 | NIST Apiezon | benzenoid | Benzene, (1-methyldecyl)- | 99 | 4536883 | C17H28 | 1143.6 | 0.02 | |
| 22.1262 | 1723 | 1735 | NIST 5%PMS VDDK | *likely contaminant* | Benzoic acid, 2-ethylhexyl ester | 96 | 5444757 | C15H22O2 | 566.2 | 0.01 | |
| 22.3029 | 1739 | 1727 | NIST HP5MS Norm | benzenoid | Benzene, (1-pentylheptyl)- | 98 | 2719622 | C18H30 | 591.0 | 0.01 | |
| 22.3534 | 1743 | 1731 | NIST HP5MS Norm | benzenoid | Benzene, (1-butyloctyl)- | 98 | 2719633 | C18H30 | 681.4 | 0.01 | |
| 22.4938 | 1756 | 1741 | NIST Apiezon | benzenoid | Benzene, (1-propylnonyl)- | 99 | 2719644 | C18H30 | 415.3 | 0.01 | |
| 22.7019 | 1775 | 1765 | NIST HP5MS VDDK | AAD | Heptadecane, 2-methyl- | 95 | 1560890 | C18H38 | 623.0 | 0.01 | |
| 23.205 | >1800 | | | | Unknown 66 | | | | 1644.3 | 0.03 | 84 (100.0%), 82 (22.7%), 67 (18.8%), 83 (13.9%), 81 (13.3%), 96 (11.3%), 96 (11.3%), 55 (11.2%), 152 (10.3%), 41 (9.3%) |
| 23.4469 | >1800 | | | | Unknown 67 | | | | 578.8 | 0.01 | 69 (100.0%), 81 (32.2%), 136 (19.7%), 41 (19.2%), 93 (16.3%), 93 (16.3%), 95 (12.5%), 57 (12.2%), 149 (10.7%), 55 (8.2%) |
| 23.6117 | >1800 | 2475 | NIST 5%PMS VDDK | *likely contaminant* | Phthalic acid, di(6-methylhept-2-yl) ester | 98 | 1000377973 | C24H38O4 | 650.8 | 0.01 | |
| 23.7442 | >1800 | 3630 | NIST 5%PMS VDDK | FAD | Methyl tetratriacontyl ether | 95 | 1000406301 | C35H72O | 855.8 | 0.01 | |
| 23.748 | >1800 | | | | Unknown 83 | | | | 370.3 | 0.01 | 68 (100.0%), 69 (49.9%), 69 (49.9%), 149 (49.3%), 83 (46.5%), 82 (43.1%), 82 (43.1%), 97 (32.9%), 111 (24.0%), 167 (15.3%) |
